# Consumption of only wild foods induces large-scale, partially-persistent alterations to the gut microbiome

**DOI:** 10.1101/2024.01.08.574648

**Authors:** Simone Rampelli, Diederik Pomstra, Monica Barone, Marco Fabbrini, Silvia Turroni, Marco Candela, Amanda G. Henry

## Abstract

The composition of the gut microbiome (GM) affects human health and varies among lifestyles. Adopting more “traditional” diets could lead to substantive and health-associated changes in the GM. However, research has focused on diets including domesticated foods. For most of our evolutionary history, humans consumed only wild foods. We explored the impact of a wild-food-only (WF) diet on the GM composition. One participant collected daily fecal samples and recorded daily food consumption over an eight-week period, the middle four weeks of which he consumed only wild foods (nuts, fruits and leafy greens, wild deer, and fish). Samples were profiled through 16S rRNA amplicon sequencing and the species identified by oligotyping. The WF diet altered the GM composition, and the magnitude of the changes is larger than in other diet interventions. However, no new GM taxa, including “old friends” appeared; instead, the relative proportions of already-present taxa shifted. There is a clear successional shift from the pre-, during- and post-WF diet. The GM is very sensitive to the change from a “Western” diet to a WF diet, likely reflecting the different macro- and micronutrient properties of the consumed foods.

## 1. Introduction

The gut microbiome (GM) is implicated in maintenance of human health [1,2]. The composition of this microbial community varies among human populations [3] and is strongly driven by differences in diet, lifestyle and living environment, and less by underlying host genetic differences [4]. More specifically, populations who consume food that is mass-produced, highly-processed and rich in fat and sugar, who have access to healthcare including antibiotics, and whose living and working spaces are often highly cleaned, have GM communities that are characterized by reduced taxonomic diversity and increased prevalence of certain taxa that are associated with inflammation and immune-mediated diseases, such as enterobacteria and mucus degraders [5]. Such populations are often labeled as having a “Western” or “industrialized” lifestyle, contrasting with “rural” or “traditional” lifestyles, which are associated with GM profiles characterized by wider biodiversity, and taxa with healthy functions, such as Prevotella and other bacteria that produce short chain fatty acids (SFCAs)[6].

Given the potential health benefits of a more diverse GM, several studies have explored how diet alterations might change GM profiles within individuals instead of among populations. Individuals accustomed to a “Western” diet and lifestyle who then relocate to an area with a more “traditional” way of life quickly acquire a more diverse GM with health-associated taxa [7]. The inverse is also true [8]. Even in the absence of a geographic change, changes to diet alone can cause significant alterations to the GM [9,10], including increasing biodiversity and the ratio of health-associated taxa. However, these diet intervention studies so far have generally remained within the realm of “Western” diets, which rely on foods from a restricted range of domesticated plants and animals. For most of the evolutionary history of our species, humans consumed only wild plant and animal foods. While the actual food items varied through time, both seasonally and over millennia, and across the many habitats in which our ancestors lived, the general composition of the diets was strikingly different from the diets consumed today, even among most “traditional” populations [11]. For example, meat from wild game has less saturated fats and more unsaturated fats than does meat from domesticated animals, while wild plants generally contain more fiber and fewer simple sugars, though overall are more variable than domesticated crops [11]. Given the link between diet and GM, and between GM and health, some have argued that returning to a diet like those of our ancestors might have health benefits. We therefore explored, in a single individual, the consequences of a wild-food-only (WF) diet on GM composition. Fecal sampling was performed daily over 8 weeks, the middle four weeks of which the participant consumed only wild food items. This is not a study of the PaleoDiet, which relies on domesticated foods and combinations of food items that would never have been available to any human ancestor in one place at one time [11]. Instead, we focused on wild (non-domesticated) foods that were available in autumn in northern Europe. Furthermore, this study isolated the effect of diet compared to other factors, such as hygiene and exposure to other people. The test subject remained living in his own house, interacting with family members, and performing other normal daily activities.

In particular, we sought to explore the following three questions. First, does adopting a WF diet influence GM composition and the prevalence of health-promoting species? Second, are there any old friend species that increase in abundance during the WF diet? Old friend species are taxa that likely have been part of the human GM in our ancestors well prior to the adoption of agriculture [12], but that are regularly not found in “Western” populations [3,13,14]. Given their long history with humans, we expect them to prefer diets like those seen in hunter-gatherer populations. Finally, do the microbiome modifications persist even after returning to a ‘normal’ diet? Previous studies indicate that some elements of the GM community reverted to the previous composition, while others remained in the altered state [2].

## 2. Materials and methods

### 2.1 Experimental design

One of us (DP) who is an experienced forager of local wild foods collected daily stool samples during an 8-week period from 2018-09-14 until 2018-11-08. The first two weeks consisted of a normal diet, followed by four weeks of a WF diet, and a further two weeks of a return to a normal diet. The wild foods were prepared using “primitive” technologies - they were cooked on an open, wood fire and processed using grindstones and flint flakes instead of modern kitchen utensils, with the exception of a meat grinder. Other aspects of the author’s lifestyle remained unchanged; he performed his usual daily activities, continued living in his house, and interacted as usual with family members. In short, this study isolated the effects of a dietary alteration. The author is of Dutch ancestry, and at the time of the experiment was 46 yo, 1.82 m tall, and weighed approximately 76 kg. The author’s weight was measured daily, and he had daily contact with a medical doctor to monitor his health and well-being during the experimental stage. All consumed food and beverage items, except water, were recorded in a daily food log. The food items in the normal diet and in the wild-food-only diet are listed in Supplementary Table 1. Ethical evaluation of the project was conducted by the Ethics Committee of the Faculties of Humanities and Archaeology at Leiden University (Letter number 2022/23).

### 2.2 Fecal collection, DNA extraction and 16S rRNA amplicon sequencing

Fecal samples were self-collected using Fe-Col® (Alpha Laboratories Ltd, Eastleigh, United Kingdom), a disposable paper device to prevent sample contamination, and SMART eNAT® (Copan SpA, Brescia, Italy) for fecal sampling and preservation. These were sent on ice to the laboratory of the Unit of Microbiome Science and Biotechnology at the University of Bologna for further analysis. All specimens were stored at -20°C until processing. Total microbial DNA for each fecal sample was extracted through a method combining bead-beating and column purification, as described previously [15].

The V3–V4 hypervariable regions of the 16S rRNA gene were amplified and library preparation was performed following the 16S Metagenomic Sequencing Library Preparation protocol (Illumina) and the Nextera technology to index libraries. Indexed libraries were pooled at an equimolar concentration of 4 nM, denatured, and diluted to 5 pM prior to sequencing on an Illumina MiSeq platform with a 2 × 250 bp paired-end protocol (Illumina, San Diego, CA, USA). Sequencing reads were deposited in the European Nucleotide Archive (ENA) with the BioProject ID XXX.

### 2.3 Bioinformatics and biostatistics analysis of GM data

All sequences were processed using a pipeline that combined PANDASeq [16] and QIIME 2 [17]. After filtering the reads by length and quality, DADA2 was used to identify the amplicon sequence variants (ASVs) [18]. Taxonomic classification was performed using the VSEARCH algorithm [19] on the SILVA database (December 2017 release) [20]. Chloroplast, mitochondria, unknown, and eukaryote sequences were removed. Oligotyping [21] was then used for clustering the high-quality filtered fasta sequences from the QIIME 2 pipeline as previously illustrated by de Goffau and colleagues [22]. In particular, we used the ‘Minimum Entropy Decomposition’ (MED) option for sensitive partitioning of high-throughput marker gene sequences from the oligotyping software with the options -M 100 (to define the minimum abundance of an oligotype) and -V 2 (to define the maximum variation allowed in each node). The final node representative sequence of each oligotype was used for species profiling using the VSEARCH algorithm and the Genomes from Earth’s Microbiomes (GEM) catalog [23] as reference database. Alpha diversity was calculated using the number of observed ASVs, the Shannon index and the Faith phylogenetic diversity index. For beta diversity, the UniFrac dissimilarities were used to construct Principal Coordinates Analysis (PCoA) plots.

Biostatistics analysis and graphical representation were performed in R using the base, vegan [24] and made4 [25] packages. Data separation in the PCoA was tested using a permutation test with pseudo-F ratios (function adonis in vegan). Kruskal-Wallis tests and Wilcoxon rank-sum test were used to assess significant differences in alpha diversity and taxon relative abundance between groups. P-values were corrected for multiple testing using the Benjamini–Hochberg procedure. A false discovery rate (FDR) ≤ 0.05 was considered statistically significant.

### 2.4 Network analysis

Species-level bacterial co-abundance groups (CAGs) were identified as previously described [3,26,27]. Briefly, the associations among taxa were determined using the Kendall correlation test, visualized with an heatmap and a hierarchical Ward-linkage clustering based on Spearman correlation distance metrics. The network plots were created using Cytoscape [28]. Circle sizes were proportional to species- or genus-level abundance or overabundance, and connections between nodes represented positive (gray) or negative (red) significant correlations. Keystone species were identified taking into account the topology of the network and the relative abundance of each taxon. Specifically, keystone nodes were identified by looking at the combination of the highest values of closeness centrality, betweenness centrality and degree on Cytoscape as previously described [29,30] and selecting only the taxa with a mean relative abundance > 1%.

### 2.5 GM across lifestyle, dietary habits and geography

GM dynamics observed in this research were compared to the results from other studies on (i) travelers in a setting with a traditional diet and lifestyle [7], (ii) people that radically changed their diet to a completely plant-based or animal-based diet [9] and (iii) Western or traditional populations [3,14,31–41].

Data from [7] and [9] were directly downloaded from the Qiita website [42], selecting the tables “55266”, “63513” and “63516”. Each table contained the OTU abundance obtained using the QIIME pipeline with the closed-reference approach and the Greengenes database (version 13_8). Only the longitudinal samples from travelers and from people that radically changed their diet to an exclusively plant-based or animal-based diet were retained. Samples from our study were reanalyzed using the same parameters reported into the Qiita website and then merged in a new table for subsequent analyses. PCoA, genus abundance superimposition to the multidimensional space and graphical representations were obtained using vegan.

For the meta-analysis using data from previous studies on subjects from different geographical locations following different subsistence strategies, we analyzed paired-end sequences using QIIME 2 [17]. The sequences were taxonomically assigned using the feature-classifier “classify-hybrid-vsearch-sklearn” option, implemented into the VSEARCH options [19] of the QIIME 2 pipeline, followed by a “q2-feature-classifier” trained on the SILVA 138.1 database [20] previously processed with RESCRIPt [43] using the developer’s instructions. The resulting abundance tables, one for each study included, were merged and rarefied, resulting in a total of 966 repository-derived samples [3,14,31–41] that were then included in the analyses along with the 57 samples generated in this work. The dataset included 407 individuals from present-day tropical and subtropical hunter-gatherer groups and 24 from Inuit tribes – most of which are undergoing a rapid transition away from their traditional hunter-gatherer diet toward a more “Western” diet – 51 individuals from rural groups practicing small-scale or subsistence agriculture from Africa, South America and Papua New Guinea, 38 individuals from Native American tribes, 12 urban Nigerians and 434 urban dwellers from North America, Europe and Asia.

The PCoA analysis to produce the beta diversity plot was performed using the vegdist function in vegan, computing Bray-Curtis distances on relative abundances at the genus level, considering only genera showing more than 0.2% of relative abundance in at least 3 samples. Compositional data were fit onto the ordination implementing the envfit function in vegan, and only genera with FDR-corrected p-values < 0.001 were plotted.

### 2.6 Functional inference of GM functions

KO (KEGG ortholog) gene abundances were predicted using the Phylogenetic Investigation of Communities by Reconstruction of Unobserved States (PICRUSt2) software [44] by applying the default parameters, including a Nearest Sequenced Taxon Index (NSTI) value of 2. Significant differences among periods are tested by Kruskall-Wallis test and represented by box plots. For beta diversity, the Bray-Curtis dissimilarity was used to build a PCoA graph and the separation verified with a permutational test with pseudo-F ratios (function adonis, in vegan).

## 3 Results

### 3.1 Individual experience of the wild-food diet

The main staples of the WF diet were chestnuts and acorns, which had to be shelled and were usually then ground to make porridge. These were supplementented by a few other nuts and seeds (hazelnuts and water lily seeds), a variety of fresh greens, dried berries and fruits, and a small amount of deer meat and ocean fish. During the WF period, DP gradually lost 4 kg, most during the first week of the WF diet. Two kg were quickly regained upon returning to a normal diet. Subjectively, DP became bored with the limited foods available to him, as there was little time to prepare more than very basic meals or to gather foods from a wider area. This likely contributed to the overall caloric reduction and weight loss. During this period, DP kept a vlog of his experiences, which is available on Youtube [45]

### 3.2 Characterization of the GM during the three dietary periods

In the pre-WF period, the GM was dominated by taxa belonging to the three major human-associated phyla, *i.e.*, Firmicutes, Bacteroidetes and Actinobacteria. In particular, the most representative families were *Lachnospiraceae*, *Ruminococcaceae*, *Bacteroidaceae*, *Oscillospiraceae*, *Rikenellaceae* and *Bifidobacteriaceae* (Fig. 1 A), which are commonly found in healthy people living a “Western” lifestyle [46]. Beta-diversity analysis revealed a clear pattern towards segregation of the microbial communities according to the sampling period, as shown by the unweighted and weighted UniFrac distances (permutation test with pseudo F-ratio, p-value ≤ 0.001) (Fig. 1 B-D). During the WF period, the GM configuration became significantly enriched in *Lachnospiraceae*, *Streptococcaceae*, *Erysipelatoclostridiaceae*, *Butyricicoccaceae*, and *Eggerthellaceae* and depleted in *Bifidobacteriaceae*, *Rikenellaceae*, *Oscillospiraceae*, *Ruminococcaceae*, *Clostridiaceae*, *Dialisteraceae*, *Acutalibacteriaceae*, and *Peptostreptococcaceae* (P< 0.05, Kruskall-Wallis test) (Fig. 1 E, see also Supplementary Table 2 for further details). Most of the modifications observed in the WF period returned to initial relative abundance values in the post-WF period, except for *Bifidobacteriaceae*, *Rikenellaceae*, *Oscillispiraceae*, and *Dialisteraceae* that remained at comparable levels to during the WF period (P < 0.05). The *Akkermansiaceae* family was even further enriched in the post-WF period compared to the two previous periods (P <0.05). In exploring differences in alpha diversity among periods, we observed a gradual increase of biodiversity from the pre-WF period, to WF and post-WF periods (P < 0.05, Kruskall Wallis test, Fig. 1 F), indicating that the intervention had an effect on the microbiome structure even after its conclusion.

**Figure 1.**
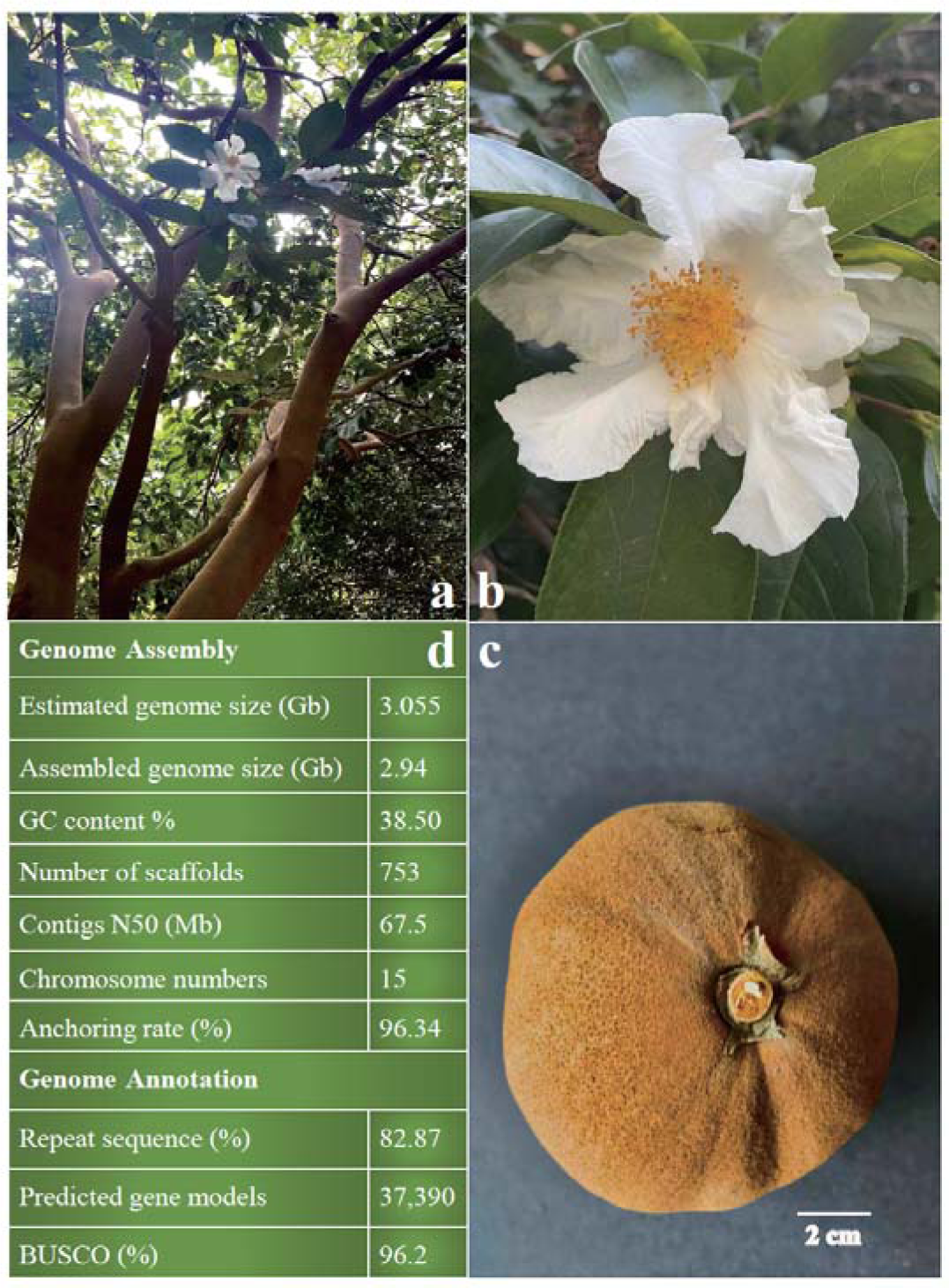
Individual experience of wild-food (WF) diet is associated with a gut microbiome shift. Comparison of microbial communities between fecal samples from pre-WF, WF and post-WF periods, represented by barplots of the family-level relative abundances (A), PCoA plots based on unweighted and weighted UniFrac distances (B,C), and boxplots for intra-group distances (D), bacterial families (E), and alpha-diversity (F). Only bacterial families that showed a significant difference in terms of relative abundance among groups are represented (P<0.05, Kruskall-Wallis test).

These changes in the proportions of individual taxa were also mirrored by changes in clusters of co-associated bacteria, which is unsurprising given the high level of interdependence within the GM. To characterize these clusters of bacteria, we generated a heatmap based on the Kendall’s tau correlation coefficients between the different 57 genera and species with a minimum relative abundance of 0.1% in at least 20% of samples. We clustered correlated bacterial species into six co-abundance groups (CAGs), indicated by different colored squares, whose relationships are represented by a Wiggum plot, where species/genus abundance is proportional to the circle diameter (Fig. 2 A and C). The dominant taxa for each CAG were *Blautia* (gray), *Streptococcus* (blue), *Coprococcus comes* (green), *Erysipelatoclostridium* (yellow), *Faecalibacterium prausnitzii* (cyan) and *Ruminococcus bicirculans* (pink). The topological data analysis indicated that *Faecalibacterium prausnitzii* and *Blautia* are the two taxa with the highest combination of 1) closeness centrality (0.46 and 0.45, respectively), 2) betweenness centrality (0.04 and 0.03, respectively) and 3) degree (15 and 11 respectively), with a mean relative abundance > 1%. For these reasons they were reported as keystone taxa for the microbial community. The CAGs changed in relative abundance across the three dietary periods (Fig. 2 B). The overabundance plots of the CAG members in the 3 dietary periods showed the emergence of different patterns of correlated microorganisms, which were found to be associated with the dietary periods (*i.e.*, pre-WF, WF and post-WF; Fig. 2 D). In particular, the GM of the pre-WF period was characterized by a *F. prausnitzii*-centered CAG, with several co-abundant glycan degraders, such as *Bacteroides* spp. (pectin, mannan, glucan, mucin) and *Bifidobacterium* (milk oligosaccharides) [47]. One auxiliary CAG was closely correlated to these bacteria and included *R. bicirculans*, *Dialister invivus*, *Bacteroides stercoris*, *Romboutsia timonensis* and CAG-83 taxon of the *Oscillospiraceae* family, which are eclectic bacteria with different substrate propensities [48–52].

**Figure 2.**
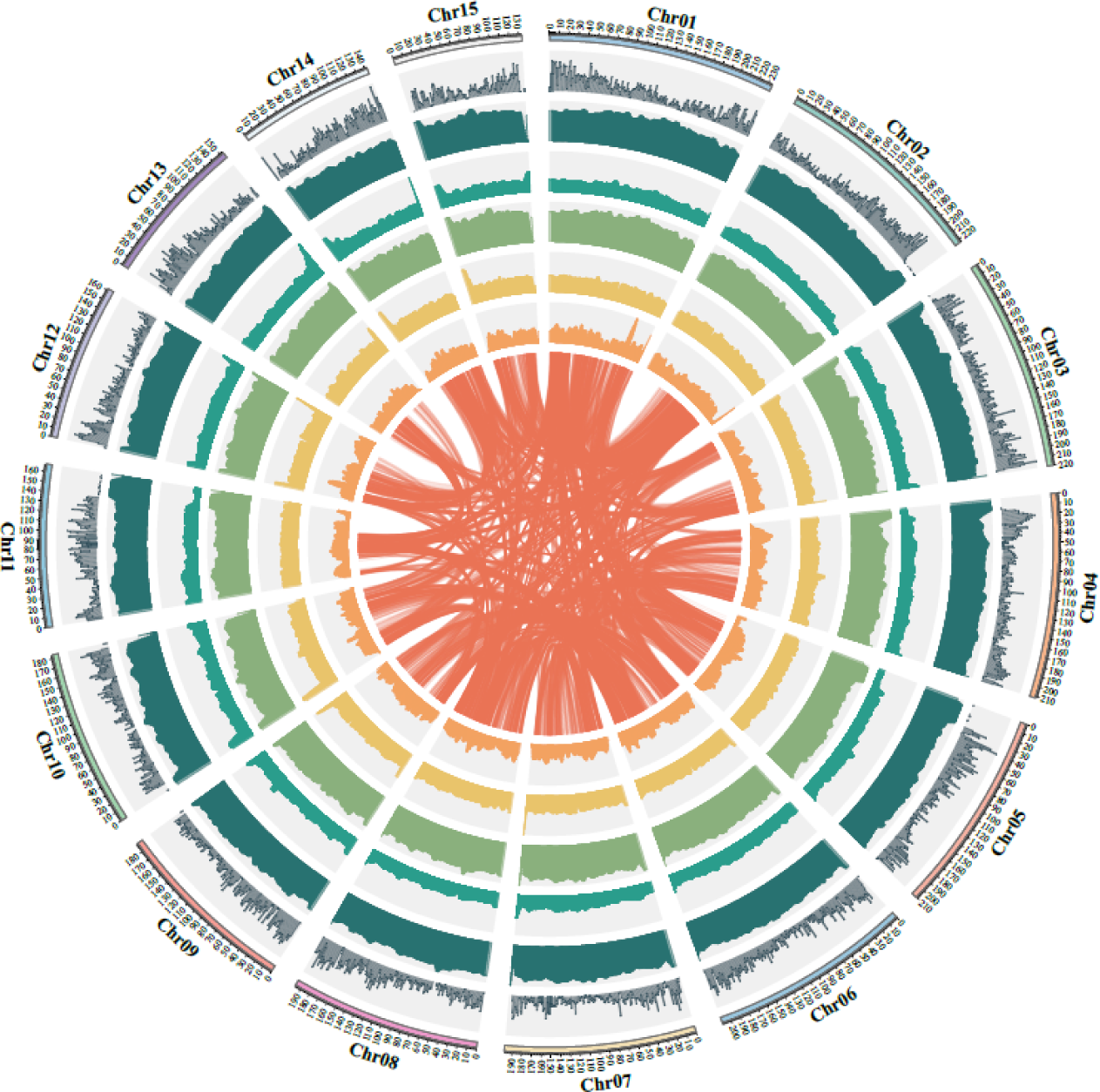
Co-abundance analysis highlights distinct bacterial networks characterizing the three dietary periods. (A) A network heatmap based on Kendall’s correlation coefficient and gut microbiome data was generated using the most abundant taxa at the genus or species level across all samples. The most dominant clusters identified are highlighted by different colored boxes and were confirmed by permutation tests with pseudo-F ratios (P<0.05, adonis of the R package vegan). One setting was used for cluster analysis (red dashed lines), which identified six clusters. The *F. prausnitzii* cluster is highlighted in cyan, the *R. bicirculans* cluster in pink, the *Erysipelatoclostridium* cluster in yellow, *C. comes* in green, *Streptococcus* in blue and *Blautia* in gray. The main representative taxa of each cluster are marked with a dot. The keystone taxa for the network structure, as highlighted by the network analysis of betweenness centrality, closeness centrality and degree, are marked with a star. The mean relative abundances for each taxon in the overall cohort are reported next to the taxon name. (B) Cumulative relative abundance of the different groups of taxa among the three periods (* P<0.001, *** P<0.00001; Kruskall-Wallis test). (C) Bacterial network scheme. Only significant Kendall’s tau correlation coefficients were considered. The leading taxa in each network are highlighted. A positive correlation is shown with a gray line and a negative correlation with a red line. Disc size is proportional to the mean relative abundance in the whole cohort. (D) Network plots corresponding to the three periods from the whole cohort analysis, in which disc sizes indicate genus/species over-abundance compared to the average relative abundance in the whole cohort.

Conversely, the WF-type GM was found to be centered around the *Blautia* CAG, which included a plethora of well-known fiber-degrading and SCFA-producing bacteria, such as taxa within the *Coprococcus eutactus* group, *Agathobaculum butyriciproducens* group and *Lachnospira rogosae* group, as well as *Blautia*, *Anaerostipes hadrus*, *Fusicatenibacter saccharivorans* and *Lachnobacterium bovis* [53,54]. Strongly associated to this cluster, the increase in members of the *C. comes* CAG and the concomitant presence of all the other CAGs enriches the WF group with a wider metabolic potential than in the previous period.

As expected, the post-WF period was characterized by an intermediate configuration between the pre-WF and the WF periods, suggesting a reapproach, although not complete, to the initial profile. Indeed, we observed a strong increase of members of the *F. prausnitzii* CAG, such as *Bacteroides* spp., to values comparable with the pre-WF period, together with the resilience of some members of the *Blautia* and *C. comes* CAGs. Notably, this period was also characterized by a higher abundance of the mucin degrader *Akkermansia muciniphila*, than the previous two periods.

Unlike the previous CAGs, the *Streptococcus* and the *Erysipelatoclostridium* CAGs did not show appreciable variations among the different dietary regimes, but only some relevant associations of specific members to each period. For instance, higher proportions of the proteolytic and animal fat-degrading taxa, such as *Alistipes shahii*, *Alistipes putredinis* and *Clostridium disporicum* were representative of the pre-WF period, whereas higher abundances of the SCFA producers *Eubacterium hallii*, *Eubacterium ramulus*, *Streptococcus* and *Erysipelatoclostridium* were characteristic of the WF period [9].

Collectively, the WF consumption caused a deep modification of the GM structure, but the GM nevertheless maintained a relevant level of inertia to partially return to the initial configuration when DP resumed a normal diet. However, there were some traits that differentiated the GM between the pre- and post-WF periods: (i) some members of the *Blautia* and *C. comes* CAGs remained at higher abundances respect to the pre-WF period; (ii) the post-WF microbiome was characterized by new traits respect to the initial period, *e.g.*, the higher abundance of *A. muciniphila*.

Putative functional changes corresponding to the observed taxonomic variations were obtained by inferred metagenomics. Given the changes in individual taxa and CAGs, some of the functional aspects of the GM were altered during the WF period as well. The GM of the WF period showed a higher propensity for starch and atrazine degradation, and phenylalanine, tyrosine and tryptophan biosynthesis, compared to the other periods, as indicated by functional assignment of GM genes using the PICRUSt2 tool (P<0.05, Kruskall-Wallis test) (Fig. 3). These changes seemed to mirror the changes to the diet that included a very heavy reliance on starch-rich nuts (acorns and chestnuts) and reduced consumption of animal products (limited meat and fish, and no poultry, dairy or eggs), which may have increased the need for amino acid biosynthesis. Phenylalanine and tyrosine are both common in milk, eggs, and some meat products, while tryptophan is common in eggs and meats. These food items were limited during the WF period. While atrazine has been banned in the EU since 2004, this herbicide is highly persistent in groundwater [55]. The location where DP acquired the wild leafy greens included field borders and previous agricultural land. Herbicide-degrading microorganisms could be possibly acquired through the ingestion of food sources endowed with these specific microbiome components, allowing their adaptation to environments under xenobiotic threat of anthropic origin [56].

**Figure 3.**
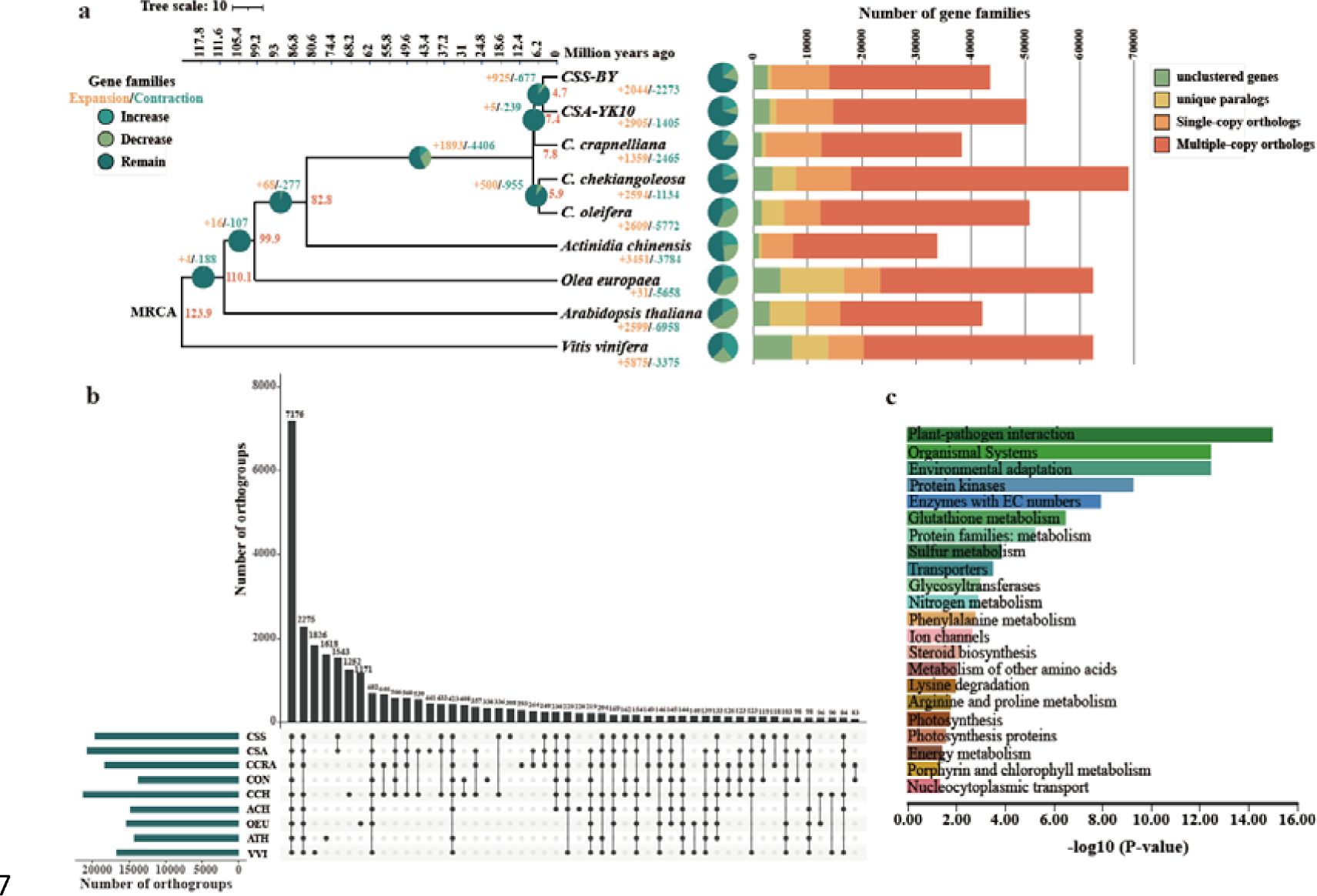
Functional characterization from inferred metagenomes allowed identifying specific traits associated with the wild food experience. (A) PCoA using Bray-Curtis distances based on functional abundance of KO genes and (B) boxplots showing significant differences (P<0.05; Kruskall-Wallis test) at pathway level of the KEGG orthology database, as inferred by PICRUSt2.

### 3.3 Diet-induced successional changes and “old friends”

The patterns observed above do not represent a stochastic change to the GM, but instead reflect a distinct successional pattern from pre-WF, to WF, to post-WF diets. The weighted proportion of species maintained, gained and lost in the temporal succession of paired samples revealed a constant ratio of species shared or newly acquired between two consecutive timepoints, while the proportion of species lost decreased slightly but significantly (P<0.05, Wilcoxon test) in the WF and post-WF periods with respect to the pre-WF period (Fig. 4). This result indicates distinct successional dynamics of the GM after the dietary modifications, with the GM keeping more diversity during and after the WF intervention. In particular, we found an increase in the number of persistent species during and after the WF period (*i.e.*, species present in at least 70% of samples for each group). The pre-WF period was characterized by the constant presence of *F. prausnitzii*, *Collinsella*, *Bacteroides vulgatus*, *Blautia*, *Gemmiger variabile* group, *Christensenella* group, *Oscillispiraceae F23-B02*, *Oscillospiraceae CAG-110*, whereas the WF period was characterized by *Blautia*, *Christensenella* group, *Oscillospiraceae F23-B02*, *Bacteroides cellulosilyticus*, *Eggerthella lenta* group, *Erysipelatoclostridium, Lachnospiraceae CAG-45*, *C. eutactus* group and *E. hallii* group. Finally, the post-WF was characterized by a higher number of persistent species, with the constant presence of *F. prausnitzii*, *Oscillospiraceae*, *Collinsella*, *B. vulgatus*, *G. variabile* group, *Blautia*, *Christensenella* group, *Dorea*, *Barnesiella intestinihominis*, *Agathobacter faecis*, *Ruminococcus bromii* group, *A. muciniphila*, *R. bicirculans* and *Adlercreutzia equolifaciens*. Notably, the list of the resilient taxa of this latter period included both some taxa characteristic of the pre-WF group, consistent with the resumption of a normal diet, but also completely new taxa that emerged after the WF diet (see the paragraph above).

**Figure 4.**
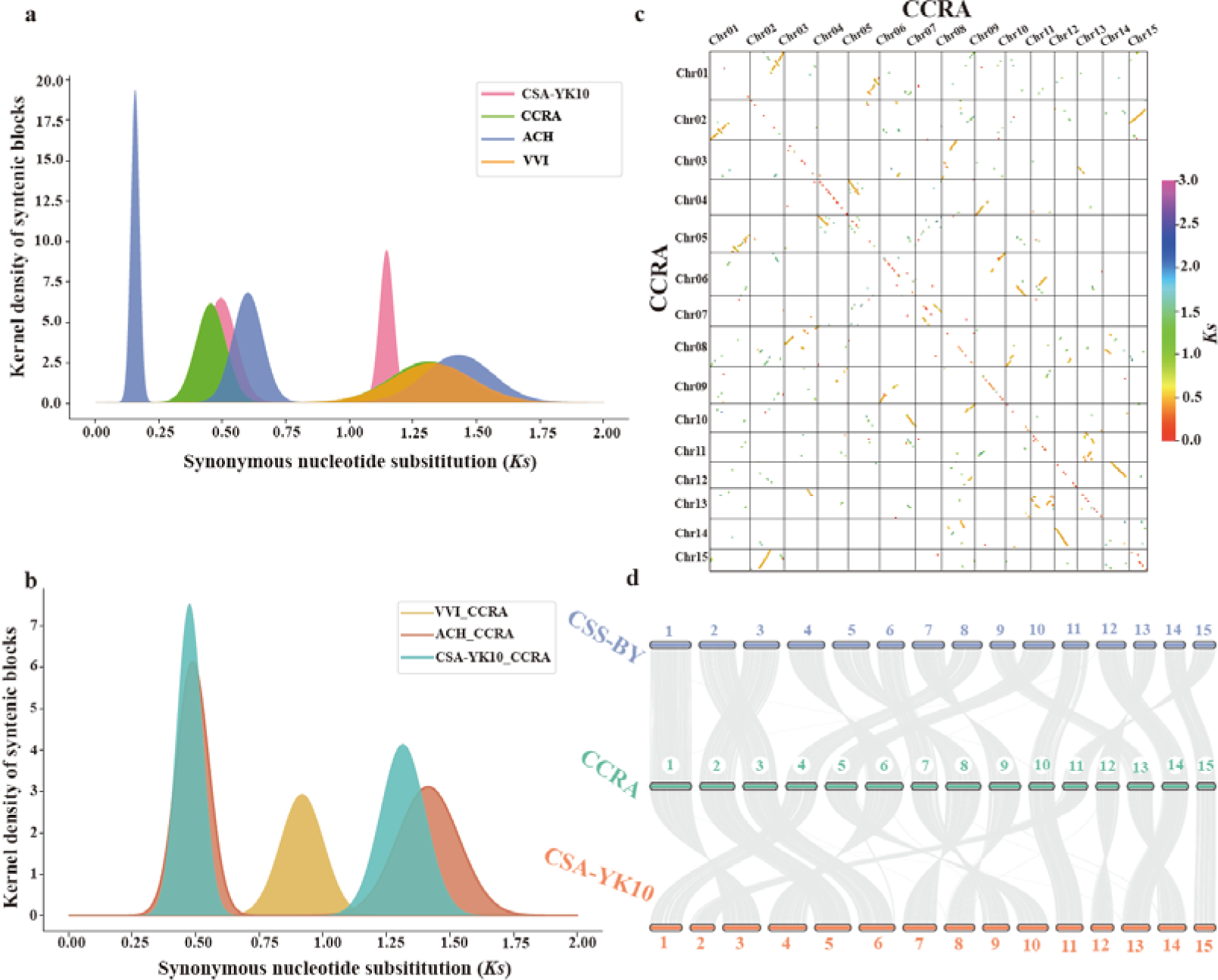
Longitudinal succession of bacterial species through the three dietary periods. (A) Weighted proportion of species maintained, gained, and lost in the downstream member of a pair of neighboring samples (* P<0.05, ** P<0.01; Kruskall-Wallis test). The black bar represents the first sample. (B) List of species belonging to the core gut microbiome of pre-WF, WF and post-WF periods. The core microbiome was defined as species present in 70% or more of samples for each group.

Despite these considerable changes to the GM communities during the dietary shifts, no old-friend taxa, *e.g.*, *Treponema*, *Prevotella* and *Succinivibrio* [3,13,14], increased during or after the WF diet. The GM changes almost exclusively involved the taxa already present into the microbial community, without the addition of new taxa. The most relevant aspect of the GM rearrangement during the transition from a normal to a WF diet was the switch of keystone species (*i.e.*, the most important taxa in defining the microbiome structure as highlighted by network analysis – see Methods for how we identified keystone taxa), from *F. prausnitzii* to *Blautia*. This supports the emerging interest in the *Blautia* genus, which has recently been proposed as a next-generation probiotic candidate, also due to its role in ameliorating inflammatory and metabolic diseases [53,57]. These changes were also associated with an overall rearrangement of butyrate-producing bacteria, from a configuration dominated by *F. prausnitzii* to a configuration where the contribution of *A. hadrus* and *E. hallii* were more relevant.

### 3.4 GM shift caused by wild-food diet is larger than in other dietary perturbations

Previous studies have explored the changes in GM structure in individuals commonly consuming a Western, industrial diet after adopting a new, more traditional diet while traveling [7], or shifting entirely to a plant-only or animal-only diet [9]. Interestingly, when compared to these, the shift in beta diversity between the initial “Western” diet and a WF-only diet was significantly larger (Fig. 5 A-C, P<0.001 permutation test with pseudo F-ratio and Kruskall-Wallis test). Furthermore, we compared the GM configurations, specifically the genus-level relative abundance profiles, of our entire study to published data from hunter-gatherer, rural agriculturalist and urban-industrial communities [3,14,31–41]. The PCoA of Bray–Curtis distances showed a clear separation between traditional and urban-industrial GMs, consistent across the different studies (Fig. 5 D). In addition, the samples from our study nested within the other urban-industrial populations, in an intermediate position between the majority of the urban-industrial samples and the GM from Native Americans and rural agriculturists. Together, our analysis highlighted how the changes in diet during the WF period instigated a rearrangement of the individual GM that did not alter in depth the microbiome structure, but acted more on the present species changing their relative abundance.

**Figure 5.**
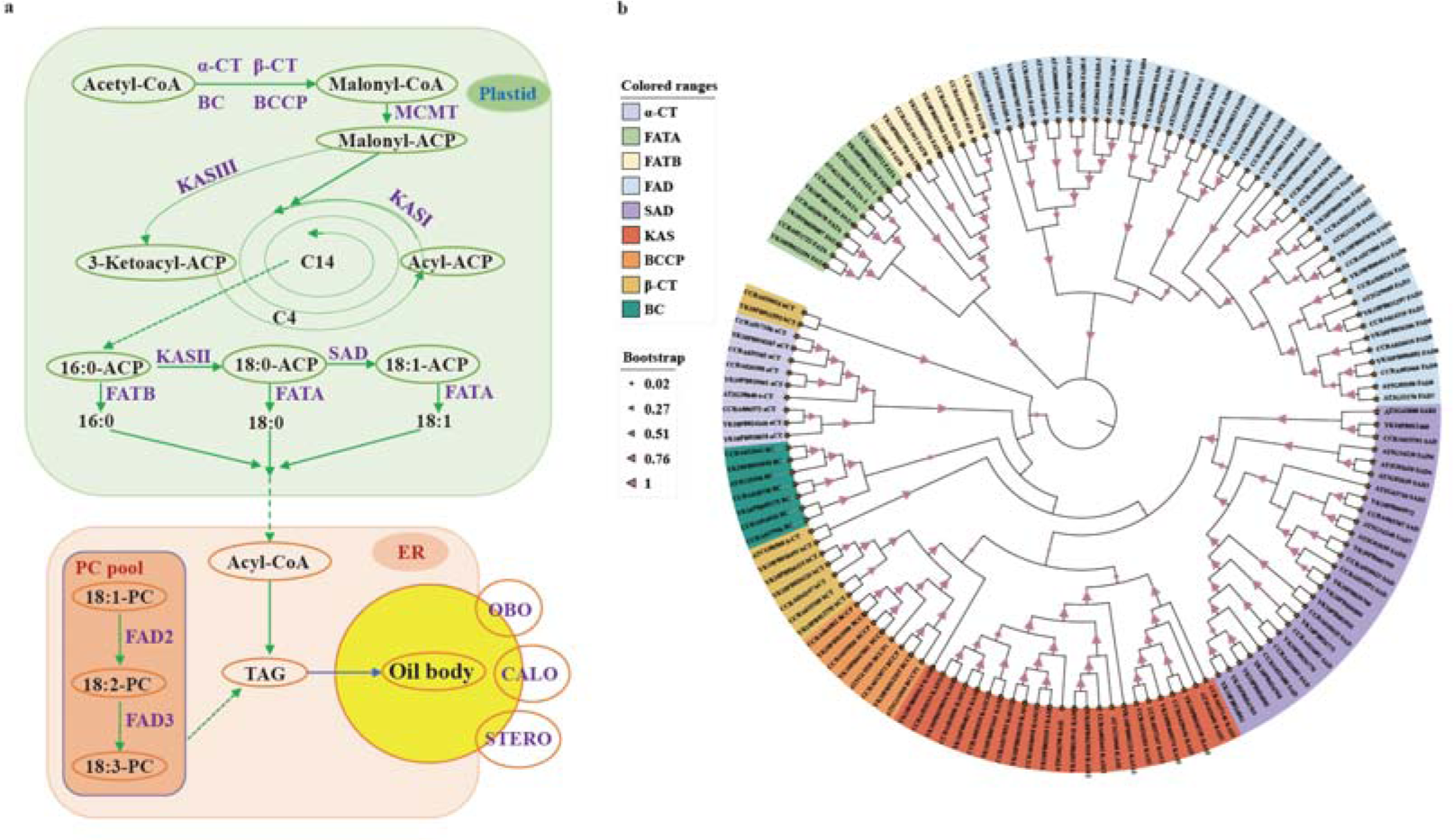
Comparison with previous studies on gut microbiome variation related to diet and lifestyle. (A) PCoA based on Bray-Curtis distances of genus-level classification using data from (i) this study, (ii) travelers in a setting with a traditional diet and lifestyle [7] and (ii) people who radically changed their diet to a completely plant-based or animal-based diet [9]. (B) PCoA coordinates on the PCo3 axis discriminated microbiome configurations of the WF period from the rest of the samples (P<0.001, Kruskall-Wallis test). (C) Superimposition of genus-level relative abundance on the same PCoA of Figure 5A reveals the most important taxa leading to the observed separation on the microbiome space (P<0.001). (D) PCoA based on Bray-Curtis distances of genus-level classification using data from this study and other works on gut microbiome characterization in populations with different geographic origin and lifestyle [3,14,31–41]. Blue arrows represent genus-level relative abundance superimposition on the PCoA space (P<0.001, permutation test).

## 4 Discussion

These results allowed us to address the three questions raised at the start of our study. First, the adoption of a diet completely lacking in domesticated foods considerably alters the human GM, and these changes are much greater than those seen in other dietary perturbations. This was surprising, given the lack of other changes to the individual’s lifestyle, including the environment and the general exposome, during this period. However, the adoption of a WF diet did not entirely alter a “Western” GM configuration to a “traditional” GM configuration, but represented a new and different arrangement of taxa. This suggests that despite considerable changes to the GM communities and relative frequencies, there remains strong inertia in the overall system, perhaps reflecting the portion of GM that is influenced by other lifestyle factors. There is some evidence for a rearrangement of health-associated taxa, including an increase in *Blautia* and the corresponding reduction in *F. prausnitzii* and *Bifidobacterium*, the latter known to be associated with the consumption of dairy foods [3]. This observation seems to corroborate the adaptive nature of the human GM, able to rearrange its compositional and functional layout keeping the homeostatic balance of the human holobiont in response to dietary shifts. Indeed, according to our findings, functional attributes of the GM community did change during the WF period, probably in response to changes in macronutrient profiles of the diet. Both the WF and the pre- and post-WF diets contained relatively small amounts of animal protein. However, the other micronutrients, including fat sources and sources of carbohydrates differed considerably, and the pre- and post-WF diets contained large amounts of dairy and eggs.

Second, despite the striking changes to the GM, there is no evidence that old friend taxa increased in abundance during the WF period. These taxa were missing in the pre-WF period and did not appear during the dietary alteration. Instead, the changes we observed were limited to abundances of already-present taxa. This suggests that WFs did not introduce new bacterial species into the GM community, despite the use of open-fire cooking and grindstones instead of modern kitchen utensils.

Third, some of the GM modifications remained even after returning to a normal diet, indicating that the adoption of a WF-only diet can induce persistent reorganization of the GM community. This altered GM profile in the post-WF period may reflect the flexibility of certain GM communities, and that there is a range of variation in taxa that are equally effective at assisting with digestion of normal diets.

Our study design is limited, focusing on one individual for only a month. Furthermore, the potential effects of the author’s change in mood during the WF diet on the GM remains unexplored. Despite this, the degree of change associated with a WF-only diet was striking in its magnitude, even when compared to other severe dietary interventions such as in [9]. This suggests that wild and domesticated foods have widely divergent properties that can be better utilized by specific GM layouts.

## List of Abbreviations

CAG: co-abundance group
GM: gut microbiome
PCoA: principal coordinate analysis
WF: wild foods

## Author Contributions

Conceptualization: AGH, DP, SR, MC; Methodology: SR, MC, DP, AGH; Investigation and Formal Analyses: SR, MB, MF, ST, MC; DNA extraction and library preparation: MB; Sequencing and revision: ST; Bioinformatic analysis: MF; Writing – Original Draft: SR, AGH; Writing – Review & Editing: SR, DP, MC, ST, AGH; Visualization: SR, MC; Funding: MC, AGH.

## Acknowledgments

This work was supported by the European Research Council under the European Union’s Horizon 2020 research and innovation program, grant agreement numbers 677576 (HARVEST: Plant foods in human evolution) and 818290 (CIRCLES: Controlling Microbiomes Circulations for Better Food Systems).

## Declaration of Competing Interest

The authors declare no conflict of interest

## Notes

### Competing Interest Statement

The authors have declared no competing interest.

## References

[1] Turroni S, Brigidi P, Cavalli A, Candela M. Microbiota–Host Transgenomic Metabolism, Bioactive Molecules from the Inside. J Med Chem 2018;61:47–61. 10.1021/acs.jmedchem.7b00244.

[2] Gilbert JA, Blaser MJ, Caporaso JG, Jansson JK, Lynch SV, Knight R. Current understanding of the human microbiome. Nat Med 2018;24:392–400. 10.1038/nm.4517.

[3] Schnorr SL, Candela M, Rampelli S, Centanni M, Consolandi C, Basaglia G, et al. Gut microbiome of the Hadza hunter-gatherers. Nat Commun 2014;5. 10.1038/ncomms4654.

[4] Vujkovic-Cvijin I, Sklar J, Jiang L, Natarajan L, Knight R, Belkaid Y. Host variables confound gut microbiota studies of human disease. Nature 2020;587:448–54. 10.1038/s41586-020-2881-9.

[5] Sonnenburg JL, Sonnenburg ED. Vulnerability of the industrialized microbiota. Science 2019;366:eaaw9255. 10.1126/science.aaw9255.

[6] Turroni S, Fiori J, Rampelli S, Schnorr SL, Consolandi C, Barone M, et al. Fecal metabolome of the Hadza hunter-gatherers: a host-microbiome integrative view. Sci Rep 2016;6:32826. 10.1038/srep32826.

[7] Ruggles KV, Wang J, Volkova A, Contreras M, Noya-Alarcon O, Lander O, et al. Changes in the Gut Microbiota of Urban Subjects during an Immersion in the Traditional Diet and Lifestyle of a Rainforest Village. MSphere 2018;3:e00193–18. 10.1128/mSphere.00193-18.

[8] Afolayan AO, Biagi E, Rampelli S, Candela M, Brigidi P, Turroni S, et al. The Gut Microbiota of an Individual Varies With Intercontinental Four-Month Stay Between Italy and Nigeria: A Pilot Study. Front Cell Infect Microbiol 2021;11:725769. 10.3389/fcimb.2021.725769.

[9] David LA, Maurice CF, Carmody RN, Gootenberg DB, Button JE, Wolfe BE, et al. Diet rapidly and reproducibly alters the human gut microbiome. Nature 2014;505:559–63. 10.1038/nature12820.

[10] Garcia-Mantrana I, Selma-Royo M, Alcantara C, Collado MC. Shifts on Gut Microbiota Associated to Mediterranean Diet Adherence and Specific Dietary Intakes on General Adult Population. Frontiers in Microbiology 2018;9.

[11] Crittenden AN, Schnorr SL. Current views on hunter-gatherer nutrition and the evolution of the human diet. American Journal of Physical Anthropology 2017;162:e23148. 10.1002/ajpa.23148.

[12] Rampelli S, Turroni S, Mallol C, Hernandez C, Galván B, Sistiaga A, et al. Components of a Neanderthal gut microbiome recovered from fecal sediments from El Salt. Commun Biol 2021;4:1–10. 10.1038/s42003-021-01689-y.

[13] Jha AR, Davenport ER, Gautam Y, Bhandari D, Tandukar S, Ng KM, et al. Gut microbiome transition across a lifestyle gradient in Himalaya. PLOS Biology 2018;16:e2005396. 10.1371/journal.pbio.2005396.

[14] Ayeni FA, Biagi E, Rampelli S, Fiori J, Soverini M, Audu HJ, et al. Infant and Adult Gut Microbiome and Metabolome in Rural Bassa and Urban Settlers from Nigeria. Cell Reports 2018;23:3056–67. 10.1016/j.celrep.2018.05.018.

[15] Viciani E, Barone M, Bongiovanni T, Quercia S, Gesu RD, Pasta G, et al. Fecal Microbiota Monitoring in Elite Soccer Players Along the 2019–2020 Competitive Season. Int J Sports Med 2022. 10.1055/a-1858-1810.

[16] Masella AP, Bartram AK, Truszkowski JM, Brown DG, Neufeld JD. PANDAseq: paired-end assembler for illumina sequences. BMC Bioinformatics 2012;13:31. 10.1186/1471-2105-13-31.

[17] Bolyen E, Rideout JR, Dillon MR, Bokulich NA, Abnet CC, Al-Ghalith GA, et al. Reproducible, interactive, scalable and extensible microbiome data science using QIIME 2. Nat Biotechnol 2019;37:852–7. 10.1038/s41587-019-0209-9.

[18] Callahan BJ, McMurdie PJ, Rosen MJ, Han AW, Johnson AJA, Holmes SP. DADA2: High-resolution sample inference from Illumina amplicon data. Nat Methods 2016;13:581–3. 10.1038/nmeth.3869.

[19] Rognes T, Flouri T, Nichols B, Quince C, Mahé F. VSEARCH: a versatile open source tool for metagenomics. PeerJ 2016;4:e2584. 10.7717/peerj.2584.

[20] Quast C, Pruesse E, Yilmaz P, Gerken J, Schweer T, Yarza P, et al. The SILVA ribosomal RNA gene database project: improved data processing and web-based tools. Nucleic Acids Research 2013;41:D590–6. 10.1093/nar/gks1219.

[21] Eren AM, Morrison HG, Lescault PJ, Reveillaud J, Vineis JH, Sogin ML. Minimum entropy decomposition: Unsupervised oligotyping for sensitive partitioning of high-throughput marker gene sequences. ISME J 2015;9:968–79. 10.1038/ismej.2014.195.

[22] de Goffau MC, Jallow AT, Sanyang C, Prentice AM, Meagher N, Price DJ, et al. Gut microbiomes from Gambian infants reveal the development of a non-industrialized Prevotella-based trophic network. Nat Microbiol 2022;7:132–44. 10.1038/s41564-021-01023-6.

[23] Nayfach S, Roux S, Seshadri R, Udwary D, Varghese N, Schulz F, et al. A genomic catalog of Earth’s microbiomes. Nat Biotechnol 2021;39:499–509. 10.1038/s41587-020-0718-6.

[24] Oksanen J, Blanchet FG, Friendly M, Kindt R, Legendre P, McGlinn D, et al. vegan: Community Ecology Package. R package version 2.5–6. 2019 2020.

[25] Culhane AC, Thioulouse J, Perrière G, Higgins DG. MADE4: an R package for multivariate analysis of gene expression data. Bioinformatics 2005;21:2789–90. 10.1093/bioinformatics/bti394.

[26] Claesson MJ, Jeffery IB, Conde S, Power SE, O’Connor EM, Cusack S, et al. Gut microbiota composition correlates with diet and health in the elderly. Nature 2012;488:178–84. 10.1038/nature11319.

[27] Rampelli S, Guenther K, Turroni S, Wolters M, Veidebaum T, Kourides Y, et al. Pre-obese children’s dysbiotic gut microbiome and unhealthy diets may predict the development of obesity. Commun Biol 2018;1:1–11. 10.1038/s42003-018-0221-5.

[28] Shannon P, Markiel A, Ozier O, Baliga NS, Wang JT, Ramage D, et al. Cytoscape: A Software Environment for Integrated Models of Biomolecular Interaction Networks. Genome Res 2003;13:2498–504. 10.1101/gr.1239303.

[29] Agler MT, Ruhe J, Kroll S, Morhenn C, Kim S-T, Weigel D, et al. Microbial Hub Taxa Link Host and Abiotic Factors to Plant Microbiome Variation. PLOS Biology 2016;14:e1002352. 10.1371/journal.pbio.1002352.

[30] Tackmann J, Matias Rodrigues JF, von Mering C. Rapid Inference of Direct Interactions in Large-Scale Ecological Networks from Heterogeneous Microbial Sequencing Data. Cell Systems 2019;9:286–296.e8. 10.1016/j.cels.2019.08.002.

[31] Clemente JC, Pehrsson EC, Blaser MJ, Sandhu K, Gao Z, Wang B, et al. The microbiome of uncontacted Amerindians. Science Advances 2015;1:e1500183. 10.1126/sciadv.1500183.

[32] Girard C, Tromas N, Amyot M, Shapiro BJ. Gut Microbiome of the Canadian Arctic Inuit. MSphere 2017;2:e00297–16. 10.1128/mSphere.00297-16.

[33] Gomez A, Petrzelkova KJ, Burns MB, Yeoman CJ, Amato KR, Vlckova K, et al. Gut Microbiome of Coexisting BaAka Pygmies and Bantu Reflects Gradients of Traditional Subsistence Patterns. Cell Reports 2016;14:2142–53. 10.1016/j.celrep.2016.02.013.

[34] Martínez I, Stegen JC, Maldonado-Gómez MX, Eren AM, Siba PM, Greenhill AR, et al. The Gut Microbiota of Rural Papua New Guineans: Composition, Diversity Patterns, and Ecological Processes. Cell Reports 2015;11:527–38. 10.1016/j.celrep.2015.03.049.

[35] Morton ER, Lynch J, Froment A, Lafosse S, Heyer E, Przeworski M, et al. Variation in Rural African Gut Microbiota Is Strongly Correlated with Colonization by Entamoeba and Subsistence. PLOS Genetics 2015;11:e1005658. 10.1371/journal.pgen.1005658.

[36] Nakayama J, Watanabe K, Jiang J, Matsuda K, Chao S-H, Haryono P, et al. Diversity in gut bacterial community of school-age children in Asia. Sci Rep 2015;5:8397. 10.1038/srep08397.

[37] Obregon-Tito AJ, Tito RY, Metcalf J, Sankaranarayanan K, Clemente JC, Ursell LK, et al. Subsistence strategies in traditional societies distinguish gut microbiomes. Nat Commun 2015;6:6505. 10.1038/ncomms7505.

[38] Sankaranarayanan K, Ozga AT, Warinner C, Tito RY, Obregon-Tito AJ, Xu J, et al. Gut Microbiome Diversity among Cheyenne and Arapaho Individuals from Western Oklahoma. Current Biology 2015;25:3161–9. 10.1016/j.cub.2015.10.060.

[39] Smits SA, Leach J, Sonnenburg ED, Gonzalez CG, Lichtman JS, Reid G, et al. Seasonal cycling in the gut microbiome of the Hadza hunter-gatherers of Tanzania. Science 2017;357:802–6. 10.1126/science.aan4834.

[40] The Human Microbiome Project Consortium. A framework for human microbiome research. Nature 2012;486:215–21. 10.1038/nature11209.

[41] Yatsunenko T, Rey FE, Manary MJ, Trehan I, Dominguez-Bello MG, Contreras M, et al. Human gut microbiome viewed across age and geography. Nature 2012;486:222–7. 10.1038/nature11053.

[42] Gonzalez A, Navas-Molina JA, Kosciolek T, McDonald D, Vázquez-Baeza Y, Ackermann G, et al. Qiita: rapid, web-enabled microbiome meta-analysis. Nat Methods 2018;15:796–8. 10.1038/s41592-018-0141-9.

[43] Robeson II MS, O’Rourke DR, Kaehler BD, Ziemski M, Dillon MR, Foster JT, et al. RESCRIPt: Reproducible sequence taxonomy reference database management. PLOS Computational Biology 2021;17:e1009581. 10.1371/journal.pcbi.1009581.

[44] Douglas GM, Maffei VJ, Zaneveld JR, Yurgel SN, Brown JR, Taylor CM, et al. PICRUSt2 for prediction of metagenome functions. Nat Biotechnol 2020;38:685–8. 10.1038/s41587-020-0548-6.

[45] Archaeologist Diederik Pomstra’s Wild Food (Palaeodiet) Experiment (part 1): introduction. 2018.

[46] Lloyd-Price J, Abu-Ali G, Huttenhower C. The healthy human microbiome. Genome Medicine 2016;8:51. 10.1186/s13073-016-0307-y.

[47] Flint HJ, Scott KP, Duncan SH, Louis P, Forano E. Microbial degradation of complex carbohydrates in the gut. Gut Microbes 2012;3:289–306. 10.4161/gmic.19897.

[48] Taylor H, Serrano-Contreras JI, McDonald JAK, Epstein J, Fell J, Seoane RC, et al. Multiomic features associated with mucosal healing and inflammation in paediatric Crohn’s disease. Alimentary Pharmacology & Therapeutics 2020;52:1491–502. 10.1111/apt.16086.

[49] Gaundal L, Myhrstad MCW, Rud I, Gjøvaag T, Byfuglien MG, Retterstøl K, et al. Gut microbiota is associated with dietary intake and metabolic markers in healthy individuals. Food Nutr Res 2022;66. 10.29219/fnr.v66.8580.

[50] Salonen A, Lahti L, Salojärvi J, Holtrop G, Korpela K, Duncan SH, et al. Impact of diet and individual variation on intestinal microbiota composition and fermentation products in obese men. ISME J 2014;8:2218–30. 10.1038/ismej.2014.63.

[51] Ricaboni D, Mailhe M, Khelaifia S, Raoult D, Million M. Romboutsia timonensis, a new species isolated from human gut. New Microbes and New Infections 2016;12:6–7. 10.1016/j.nmni.2016.04.001.

[52] Atzeni A, Bastiaanssen TFS, Cryan JF, Tinahones FJ, Vioque J, Corella D, et al. Taxonomic and Functional Fecal Microbiota Signatures Associated With Insulin Resistance in Non-Diabetic Subjects With Overweight/Obesity Within the Frame of the PREDIMED-Plus Study. Frontiers in Endocrinology 2022;13.

[53] Liu X, Mao B, Gu J, Wu J, Cui S, Wang G, et al. Blautia—a new functional genus with potential probiotic properties? Gut Microbes 2021;13:1875796. 10.1080/19490976.2021.1875796.

[54] Vacca M, Celano G, Calabrese FM, Portincasa P, Gobbetti M, De Angelis M. The Controversial Role of Human Gut Lachnospiraceae. Microorganisms 2020;8:573. 10.3390/microorganisms8040573.

[55] Jablonowski ND, Schäffer A, Burauel P. Still present after all these years: persistence plus potential toxicity raise questions about the use of atrazine. Environ Sci Pollut Res 2011;18:328–31. 10.1007/s11356-010-0431-y.

[56] Henry LP, Bruijning M, Forsberg SKG, Ayroles JF. The microbiome extends host evolutionary potential. Nat Commun 2021;12:5141. 10.1038/s41467-021-25315-x.

[57] Liu X, Guo W, Cui S, Tang X, Zhao J, Zhang H, et al. A Comprehensive Assessment of the Safety of Blautia producta DSM 2950. Microorganisms 2021;9:908. 10.3390/microorganisms9050908.

